## Supplemental Figures and Tables for "Inducible cell-specific mouse models for paired epigenetic and transcriptomic studies of microglia and astroglia"

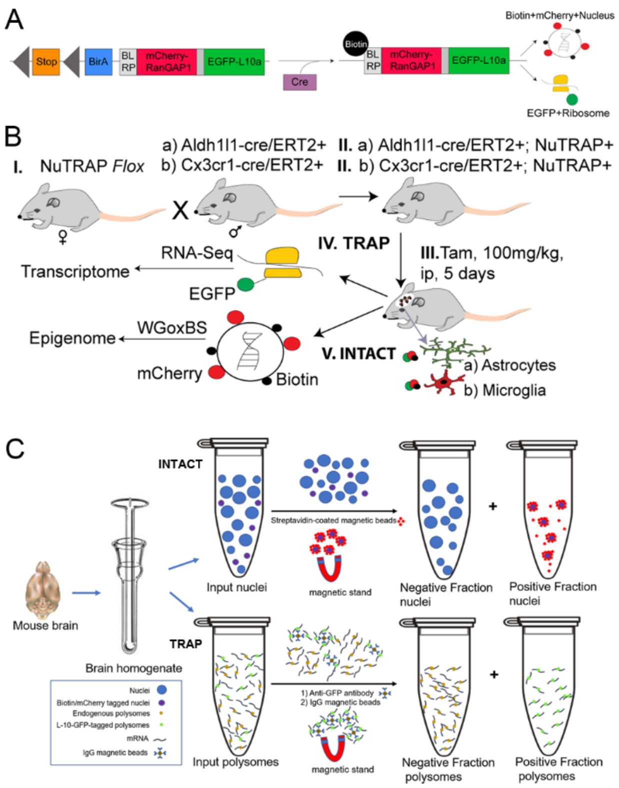


**Supplemental Figure 1. Main experimental design in the study.** A) Representation of the NuTRAP construct and its recombined products upon cre-mediated induction. B) Breeding strategy to achieve astrocyte- and microglia- specific RNA and DNA. C) Schematic of the cell-specific RNA and DNA isolation from the same mouse brain via the TRAP and INTACT methods.

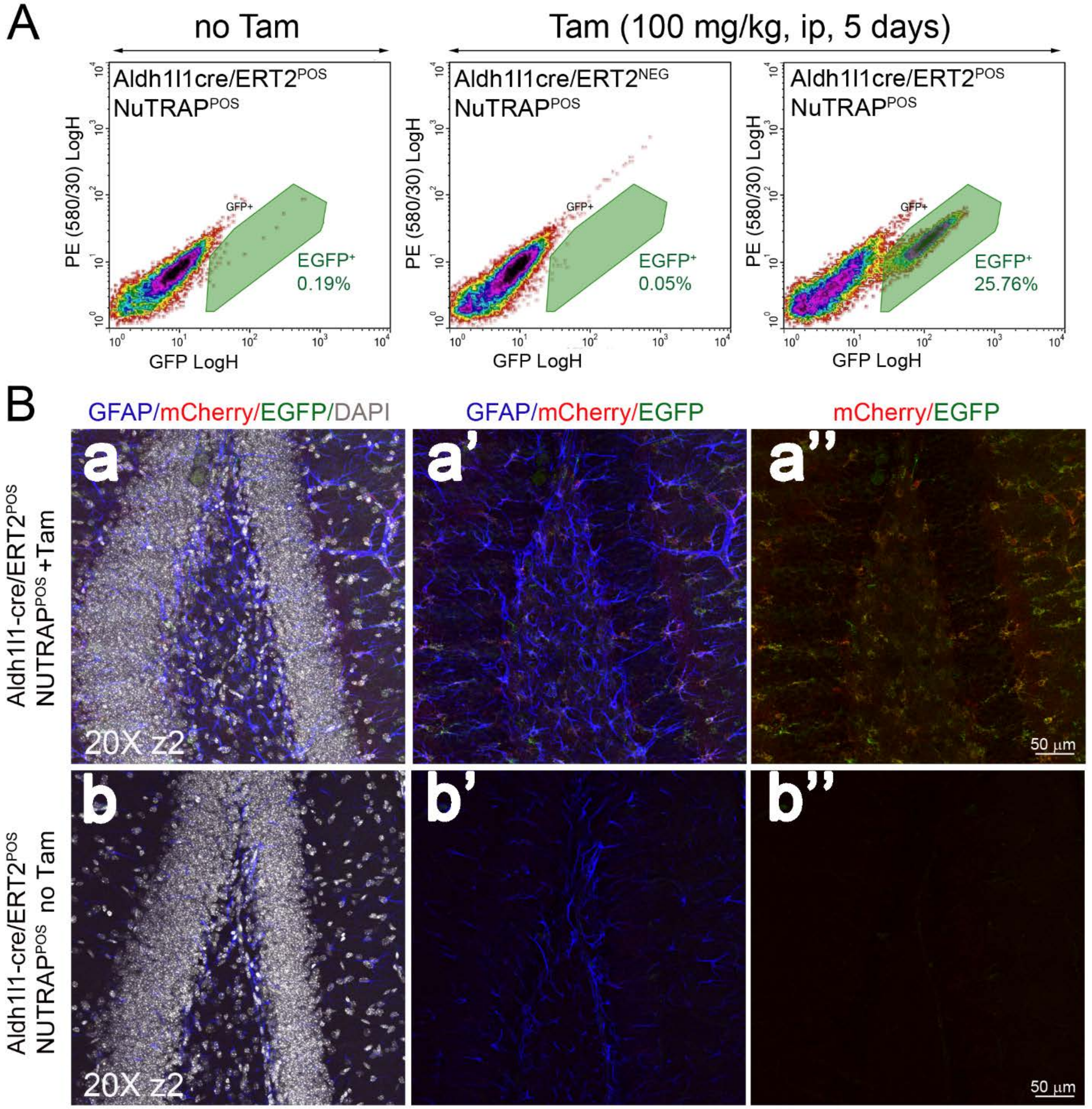


**Supplemental Figure 2. Flow cytometry and immunohistochemical assessment of Tam-independent recombination in the Aldh1l1-NuTRAP brain.** Aldh1l1-creNuTRAP and cre negative NuTRAP$^{+}$ mice were treated with Tam or left untreated and after a week brains were dissected for flow cytometry (FC) and immunohistochemistry (IHC) analyses. A) Representative FC plots of single- cell suspensions show a distinct population of EGFP$^{+}$ cells (25.76%) in brain samples of Aldh1l1-NuTRAP mice treated with Tam. Such EGFP$^{+}$ cell population was negligible (0.05-0.19%) in brains of Aldh1l1-NuTRAP mice left untreated or cre negative NuTRAP positive mice treated with Tam (n=2/group). B) Representative confocal fluorescent microscopy images of sagittal brain sections show colocalization of EGFP (green signal) and mCherry (red signal) to GFAP expressing cells (blue signal: astrocytes) in the Aldh1l1-NuTRAP. Untreated counterparts did not display EGFP or mCherry expression (n=3/group).

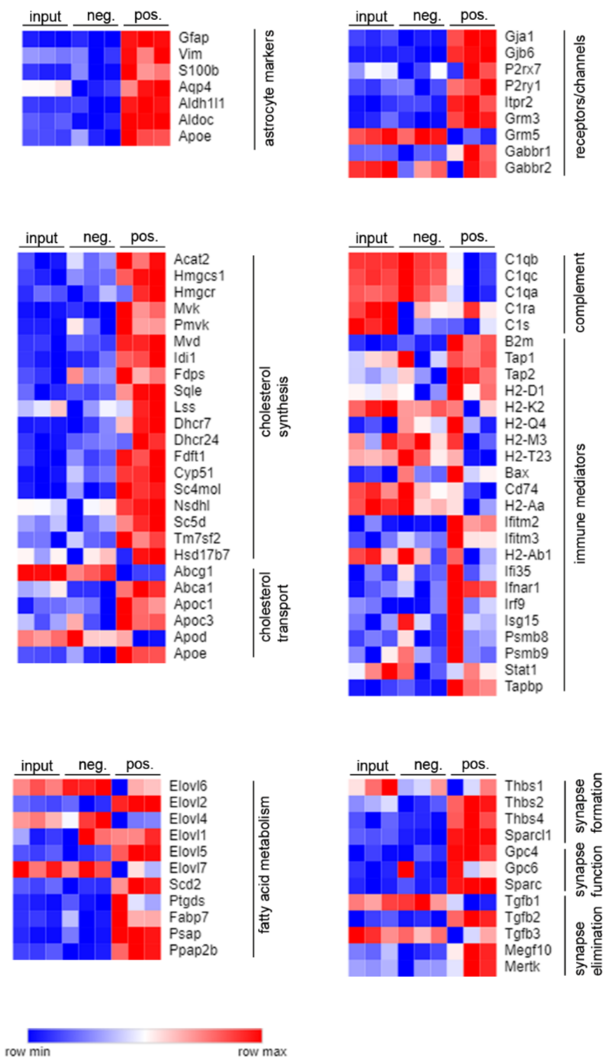


**Supplemental Figure 3. Differential gene expressions in Aldh1l1-NuTRAP positive fraction strongly correlate with physiological functions of astrocytes in the brain.** Heatmaps show in the TRAP positive fraction of Aldh1l1-NuTRAP brains is enriched in astrocyte marker genes compared to input, in agreement with over-representation of genes that are critical in astrocyte physiological functions such as cholesterol synthesis and transport, fatty acid metabolism, receptors/channels, some complement/immune mediators, and synapse modification (formation, function, and elimination) processes.

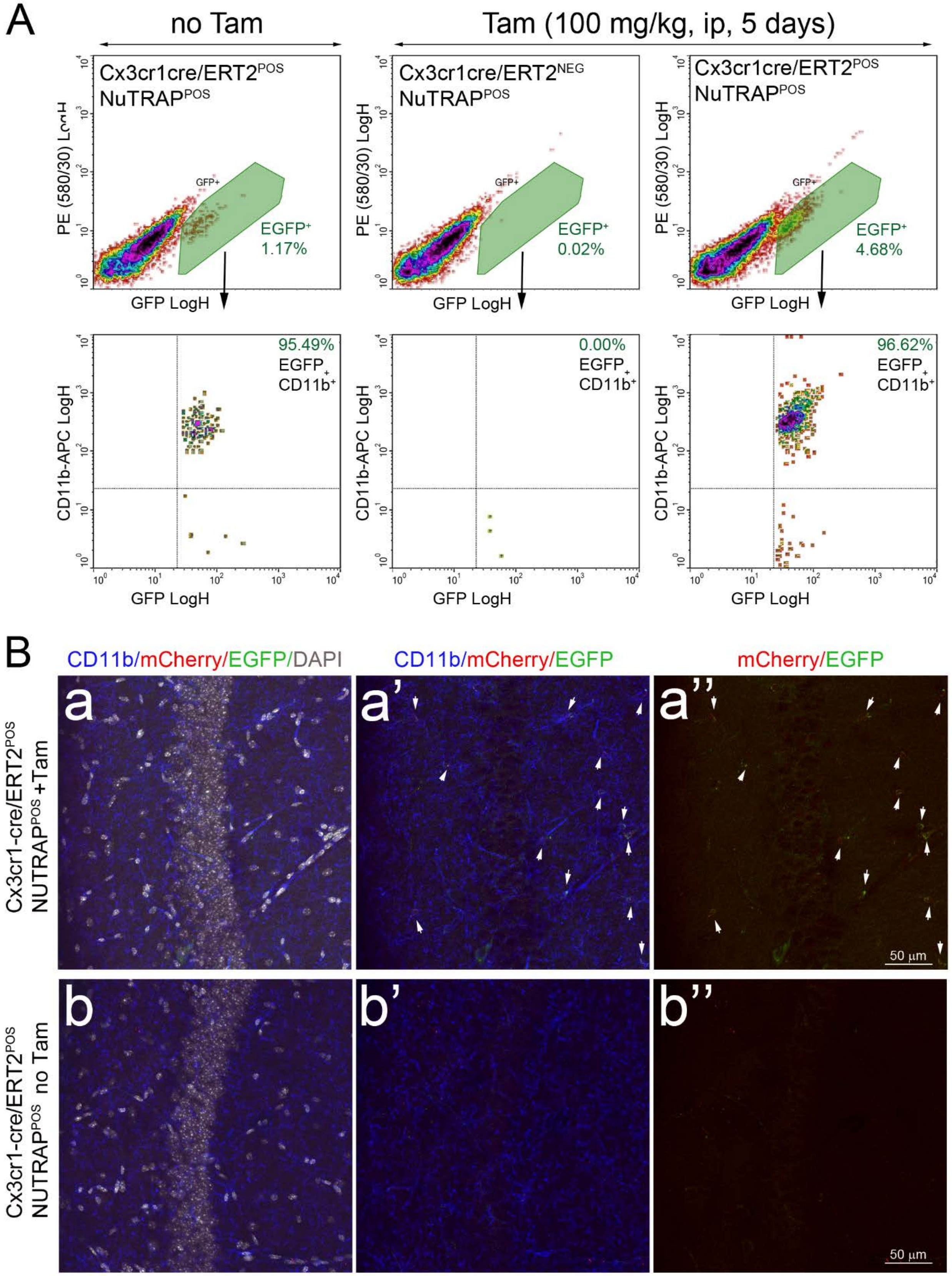
A
no Tam
Tam (100 mg/kg, ip, 5 days)
Cx3cr1cre/ERT2POS NuTRAPPOS
Cx3cr1cre/ERT2NEG NuTRAPPOS
Cx3cr1cre/ERT2POS NuTRAPPOS
PE (580/30) LogH
GFP LogH
EGFP+ 1.17%
EGFP+ 0.02%
EGFP+ 4.68%
CD11b-APC LogH
95.49% EGFP+ CD11b+
0.00% EGFP+ CD11b+
96.62% EGFP+ CD11b+
B
CD11b/mCherry/EGFP/DAPI
CD11b/mCherry/EGFP
mCherry/EGFP
Cx3cr1-cre/ERT2POS NUTRAPPOS +Tam
Cx3cr1-cre/ERT2POS NUTRAPPOS no Tam
a
a'
a''
b
b'
b''
50 µm
50 µm

**Supplemental Figure 4. Flow cytometry and immunohistochemical assessment of Tam-independent recombination in the Cx3cr1cre-NuTRAP brain.** Cx3cr1-cre/ERT2$^+$; NuTRAP$^+$ and Cx3cr1-cre/ERT2$^-$; NuTRAP$^+$ mice were treated with Tam or left untreated and after three weeks brains were dissected for flow cytometry (FC) and immunohistochemistry (IHC) purposes. A) Representative FC plots of single- cell suspensions show a distinct population of EGFP$^+$ cells (4.68%) in brain samples of Cx3cr1-cre/ERT2$^+$ mice treated with Tam that almost exclusively co-expressed CD11b (96.62%) and was undetectable (0.02 %) in brains of Cx3cr1-cre/ERT2$^-$ held to the same treatment. In Cx3cr1-cre/ERT2$^+$ mice that did not receive Tam, EGFP+ were detected at a lesser level than in treated counterparts (1.17%) (n=2/group). B) Representative confocal fluorescent microscopy images of sagittal brain sections show colocalization of EGFP (green signal) and mCherry (red signal) to CD11b expressing cells (blue signal) in the Aldh1l1-cre/ERT2$^+$. Untreated counterparts had almost no EGFP or mCherry expression (n=3/group).

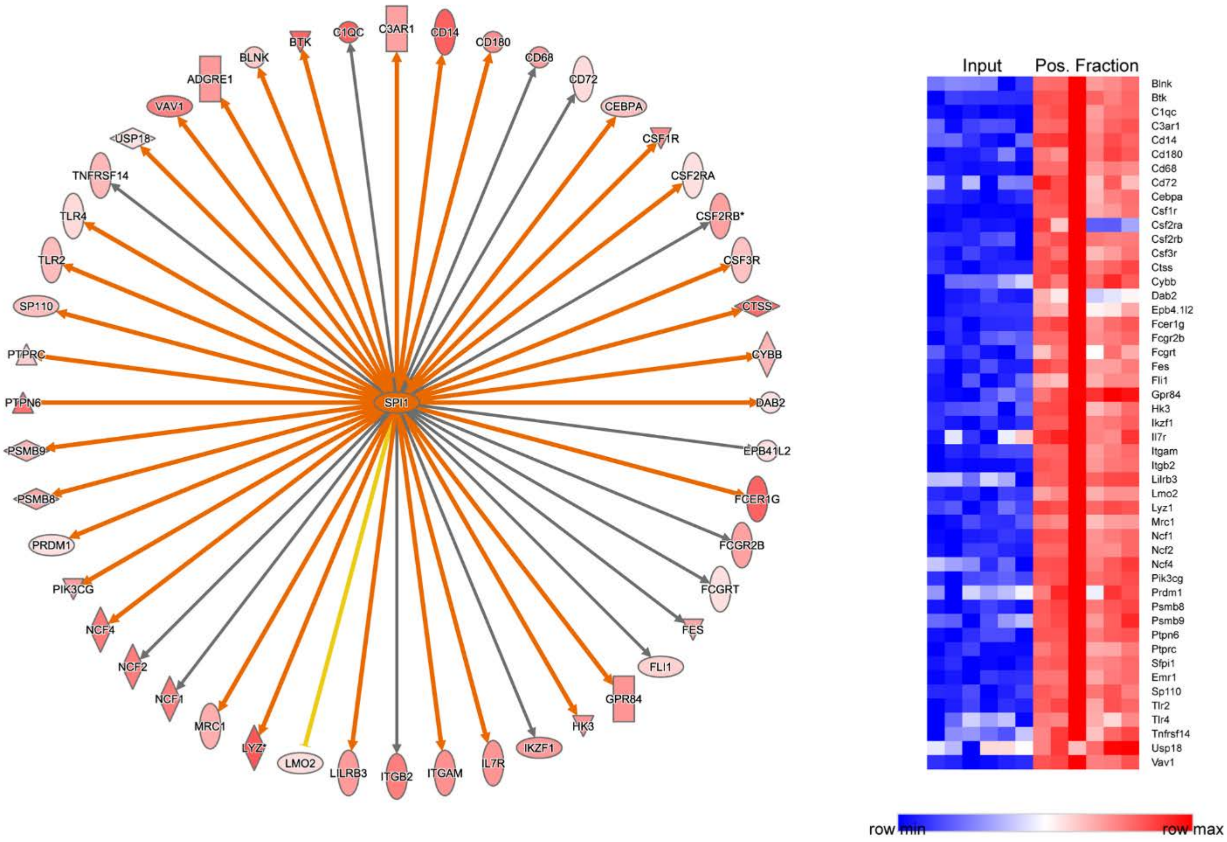


**Supplemental Figure 5. Differential gene expressions in Cx3cr1-NuTRAP positive fraction correlate with enrichment of canonical targets of microglial SPI1 in the brain.** Genes enriched in the microglia transcriptome included an overrepresentation of genes regulated by PU.1 (also known as Spi1), a transcription factor that shapes the homeostatic functions of microglia (2000-2019 Qiagen. All rights reserved).

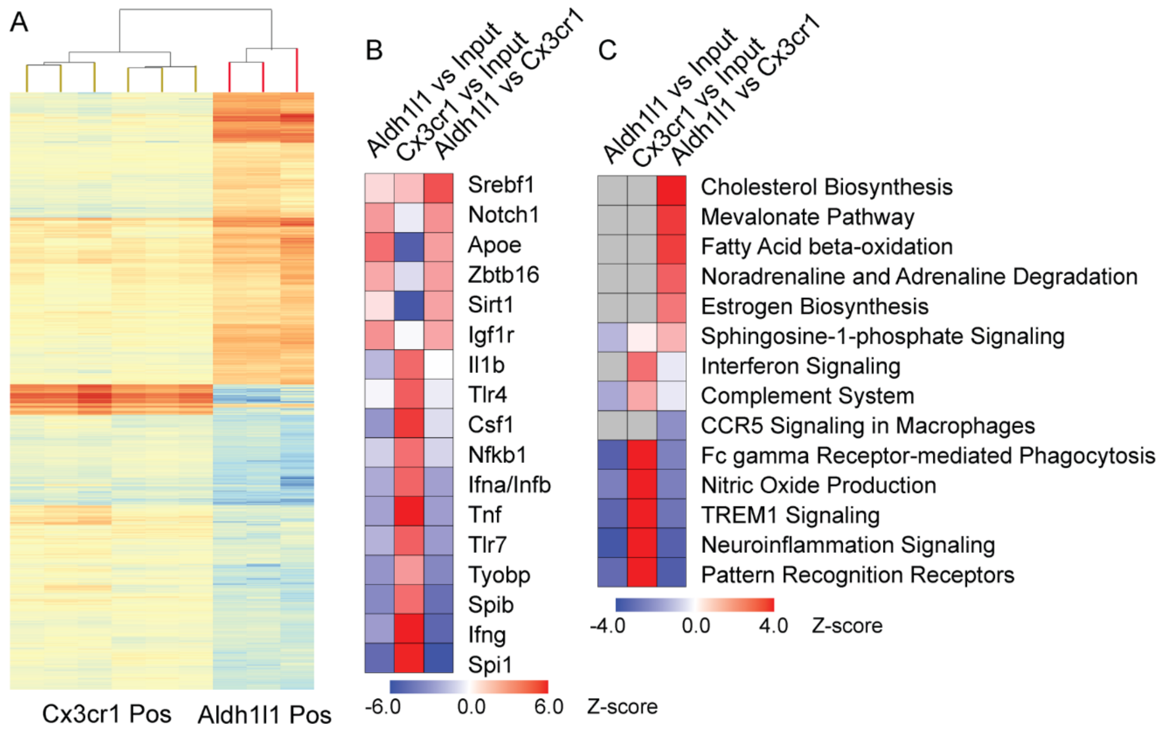


**Supplemental Figure 6. Transcriptomic comparison of astrocytes and microglia.** A) Positive fractions from Aldh1l1-NuTRAP and Cx3cr1-NuTRAP were compared. B) Regulator analysis comparison. C) Pathway analysis comparison.

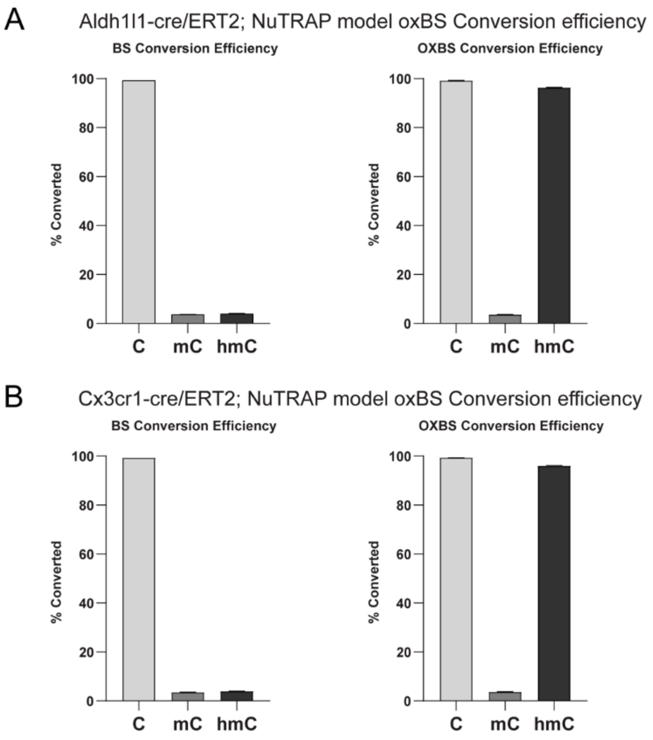


**Supplemental Figure 7. Conversion efficiency of CEGX spike-in controls.** Exogenous control sequences (CEGX, Cambridge, UK) with methylation and hydroxymethylation at specific bases were spiked in to each sheared DNA sample (0.04% w/w) prior to oxidation and/or bisulfite conversion. After sequencing, raw fastq files were run through CEGXQC v0.2, a custom FASTQC program, to generate summary documents and QC reports based on the conversion performance of the spike-in sequencing controls. Conversion percentages for different cytosine modifications (C, mC, and hmC) are plotted for bisulfite-converted and oxidative bisulfite-converted libraries. Bisulfite-converted libraries show high conversion of unmodified cytosines and low conversion of methylated and hydroxymethylated cytosines. While oxidative bisulfite-converted libraries show high conversion of unmodified cytosines and hydroxymethylated cytosines, and low conversion of methylated cytosines. The bisulfite-converted libraries are used to determine the total percent modified cytosines (mC+hmC), while the oxidative bisulfite-converted libraries are used to determine the percent methylated cytosines (mC).

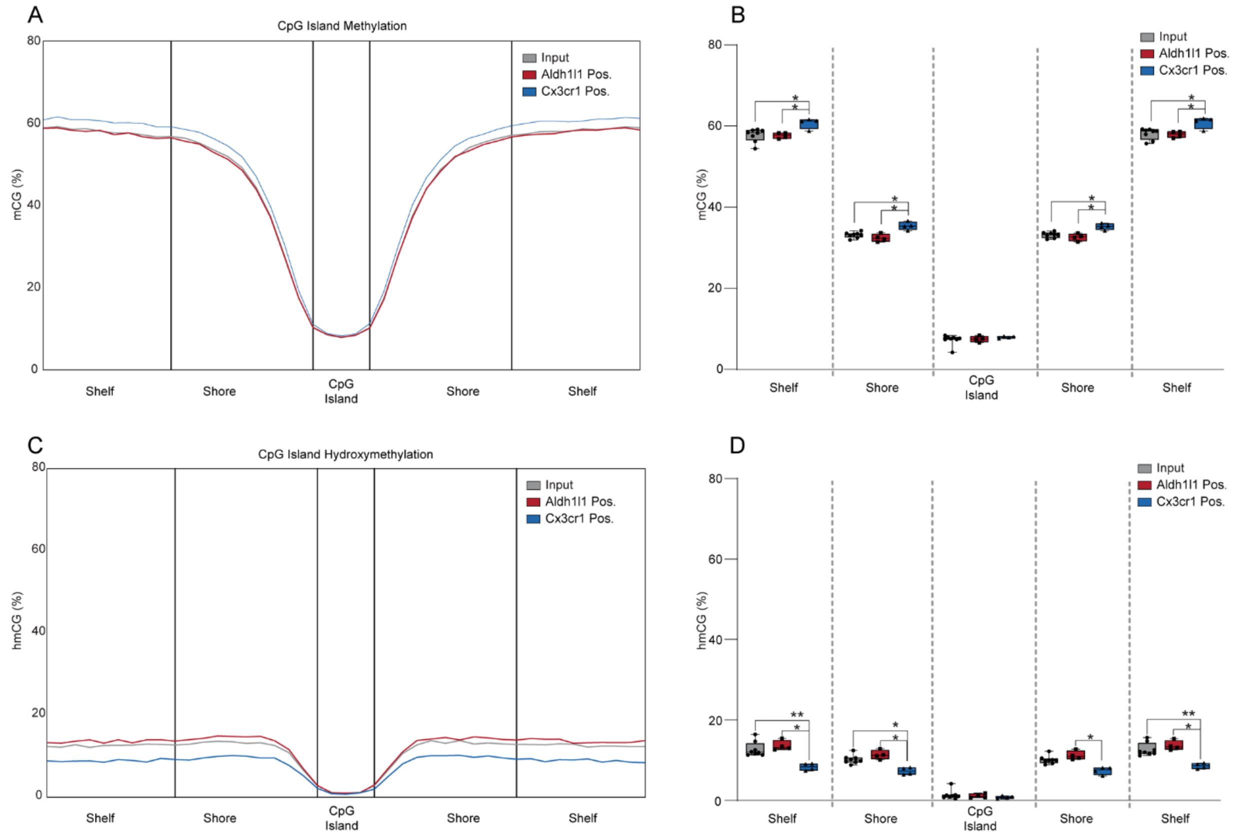


**Supplemental Figure 8. CpG island, shore and shelf methylation and hydroxymethylation in the Aldh1l1- NuTRAP and Cx3cr1- NuTRAP mouse brains by WGoxBS.** After Tam treatment, half brain hemispheres were harvested from Aldh1l1-NuTRAP and Cx3cr1-NuTRAP mice and subjected to nuclei isolation and subsequent INTACT protocol for genomic DNA extraction for epigenome analyses. Analysis of methylation and hydroxymethylation levels covering CpG islands, shores, and shelves revealed that the shores and shelves of Cx3cr1-NuTRAP INTACT positive fractions cells had significantly higher mCG levels (A-B) and significantly lower hmCG levels (C-D) compared to the other groups. (n=8/input group, n=4/positive fraction group; 2-way ANOVA with Tukey's multiple comparison test, $*p<0.05$, $**p<0.01$).

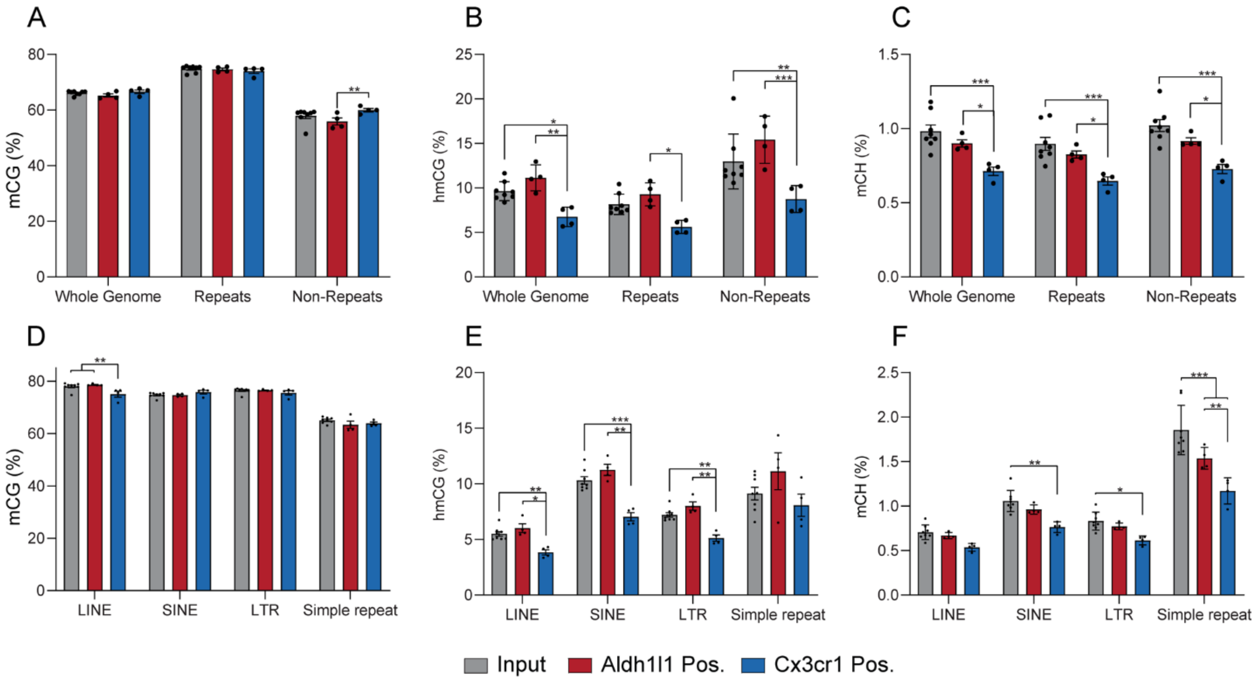


**Supplemental Figure 9. Comparison of whole genome, repeat, and non-repeat levels of methylation and hydroxymethylation between input and positive fractions of INTACT-isolated DNA from Aldh1l1-NuTRAP and Cx3cr1-NuTRAP mouse brains.** A) Overall repetitive elements have higher levels of CG methylation and non-repetitive elements have lower levels of CG methylation than whole genome levels. While there are no differences in total mCG between the input DNA, Aldh1l1+ DNA, and Cx3cr1+ DNA in the whole genome or repeat elements, there is a small, but significant, difference in mCG between Aldh1l1$^{+}$ DNA and Cx3cr1$^{+}$ DNA in non-repeat elements of the genome. B) There is less hmCG of Cx3cr1+ DNA than both input DNA and Aldh1l1+ DNA at the whole genome level and in repetitive and non-repetitive elements. C) There is less mCH of Cx3cr1+ DNA than both input DNA and Aldh1l1+ DNA at the whole genome level and in repetitive and non-repetitive elements. D) LINEs contain less mCG in Cx3cr1+ DNA than in input DNA or Aldh1l1+ DNA. (E) LINEs, SINEs, and LTRs have lower hmCG levels in Cx3cr1+ DNA than in input DNA or Aldh1l1+ DNA**.** F) SINEs and LTRs have lower mCH levels in Cx3cr1+ DNA than input DNA. Simple repeats have lower mCH levels in Cx3cr1+ DNA than in input DNA and Aldh1l1+ DNA. Aldh1l1+ DNA has lower mCH levels in simple repeats than input DNA. (n=8/input group, n=4/positive fraction group; 2-way ANOVA with Tukey's multiple comparison's test, *p<0.05, **p<0.01, ***p<0.001).

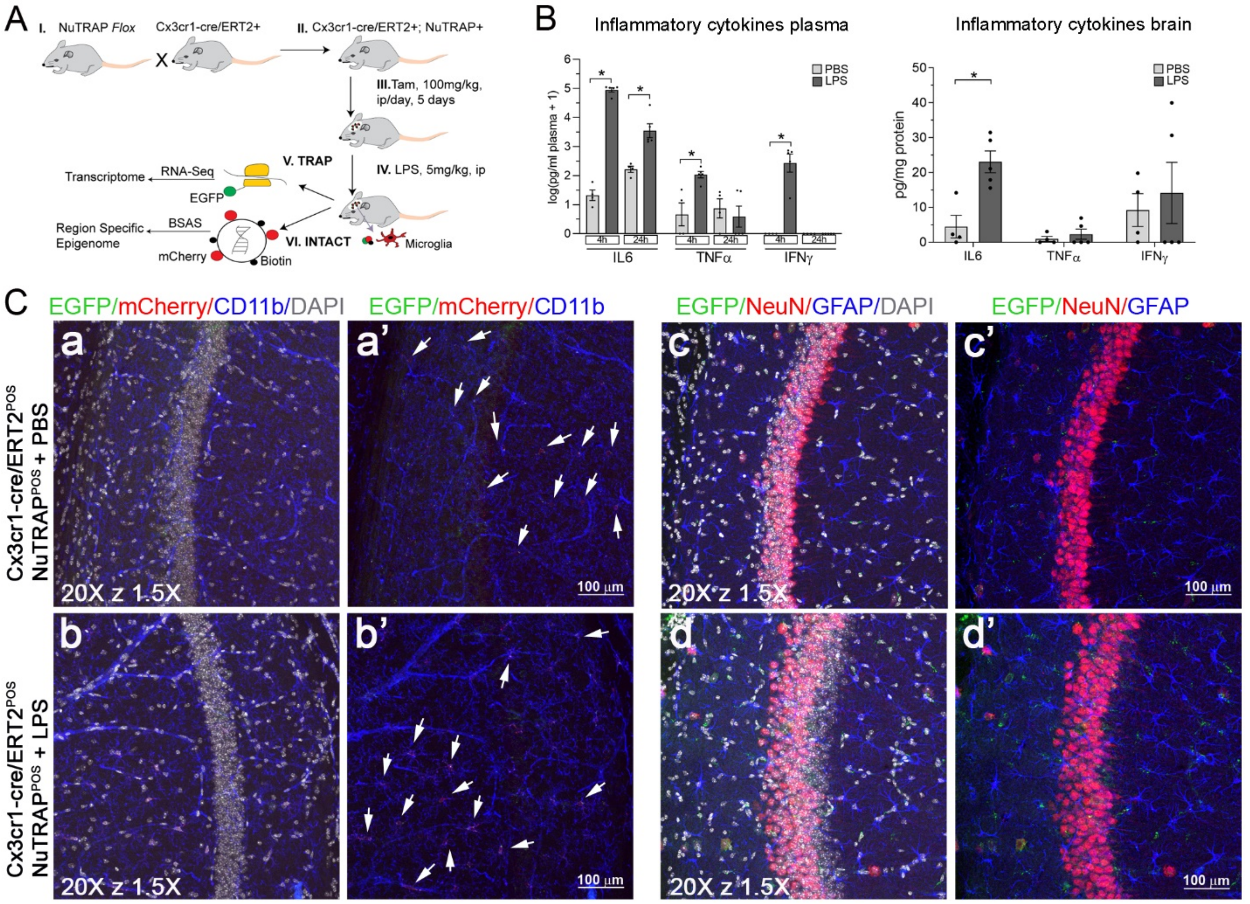


**Supplemental Figure 10. LPS systemic treatment as experimental demonstration of the Cx3cr1-cre/ERT2 model.** A) Schematic of the experimental design for epigenetic and transcriptomic analyses of brain microglia upon LPS challenge. 3-4 weeks after Tam treatment, Cx3cr1-cre/ERT2$^{+}$; NuTRAP$^{+}$ mice were subjected to a single ip injection with LPS or PBS as control and 24 h later their brains dissected for protein and IHC purposes. B) Validation of systemic LPS treatment. Blood samples were collected at 4 and 24h after LPS injection before euthanasia and brain harvest. Plasma samples and brain tissue homogenates were used to measure concentration of inflammatory cytokines IL-6, TNFα, and IFNɣ by suspension array. Values are expressed as average pg analyte/ml ± SEM in plasma (4h and 24h time points) and average pg analyte/mg ±SEM in tissue (24h time point). C) Representative confocal fluorescent microscopy images of sagittal brain sections EGFP expression (green signal) was found in cells that co-expressed mCherry (red signal) and CD11b (blue signal). * $p<0.05$ between PBS and LPS treated groups for each analyte by unpaired T test (n=4/PBS group and 5/LPS group).

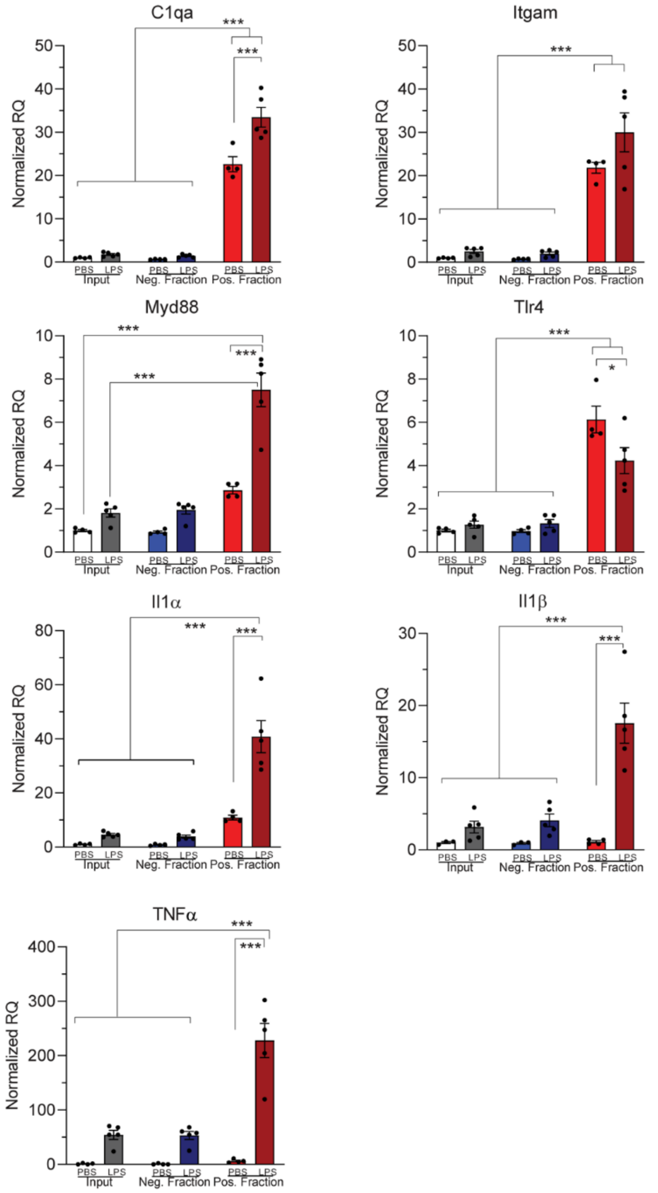


**Supplemental Figure 11. qPCR validation of TRAP-RNA in the Cx3cr1-cre/ERT2 model following LPS treatment.** 3-4 weeks after Tam treatment, Cx3cr1-cre/ERT2$^{+}$; NuTRAP$^{+}$ mice were subjected to a single ip injection with LPS or PBS as control and 24 h later their brains harvested and one hemisected half used for TRAP isolation of RNA and downstream analyses. qPCR analysis of microglial genes (C1qa and Itgam) and candidate genes related to LPS-induced inflammation (Tlr4, Myd88, Il1α, Il1β, and Tnfα) demonstrate both enrichment and higher magnitude changes in the positive fraction as compared to the input. Bar graphs represent average RQ ± SEM for each gene expression measured. *, *** $p<0.05$ and $p<0.001$ respectively by two-way ANOVA followed by the Sidak's multiple comparison test (n=4 for PBS group and n=5 for LPS group).

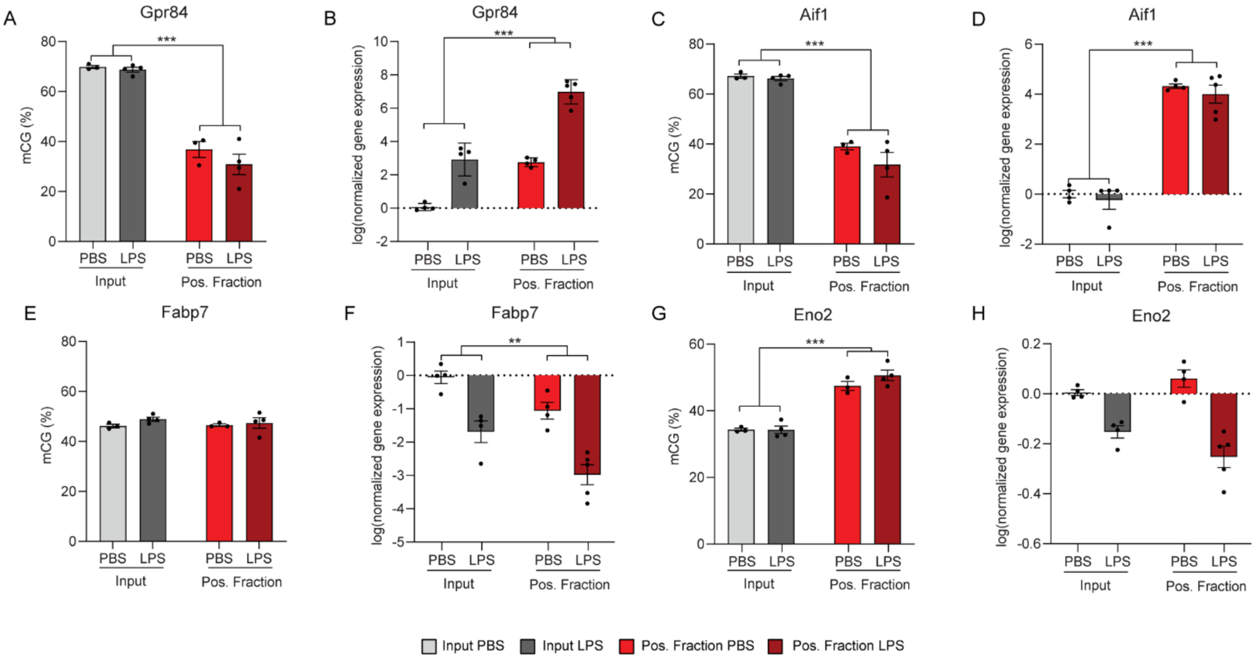


**Supplemental Figure 12. Targeted bisulfite amplicon sequencing (BSAS) to assess methylation (mCG) of microglial DNA in cell-specific marker gene promoters 24 hours after LPS challenge in Cx3cr1-NuTRAP mouse brain.** Gpr84, a microglial marker gene, has (A) lower CG promoter methylation, by BSAS, and (B) higher gene expression, by RNA-Seq, in the INTACT-isolated positive fraction than the input, regardless of LPS treatment. Aif1, a microglial marker gene, has (C) lower CG promoter methylation, by BSAS, and (D) higher gene expression, by RNA-Seq, in the INTACT-isolated positive fraction than the input, regardless of LPS treatment. Fabp7, an astrocyte marker gene, has (E) no difference in promoter CG methylation between input and INTACT-isolated positive fraction, by BSAS, and (F) higher gene expression, by RNA-Seq, in the INTACT-isolated positive fraction than the input, regardless of LPS treatment. Eno2, a neuronal marker gene, has (A) higher CG promoter methylation, by BSAS, and (B) no difference in gene expression, by RNA-Seq, in the INTACT-isolated positive fraction than the input, regardless of LPS treatment. (2-way ANOVA or mixed effects analysis with Holm-Sidak correction for multiple comparisons, main effect $*p<0.05$, $**p<0.01$, and $***p<0.001$).

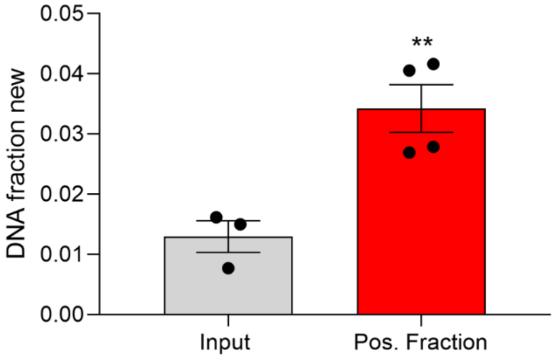


**Supplemental Figure 13. Stable isotope labeling study in the Cx3cr1-NuTRAP mouse brain.** Microglial proliferation was measured as incorporation of deuterium into purine deoxyribose. Mice were given an intraperitoneal injection of 99.9% $D_2O$ and subsequently provided drinking water enriched with 8% $D_2O$ for 30 days. Following INTACT-DNA isolation, DNA was hydrolyzed for analysis of the pentafluorobenzyl-*N*,N-di(pentafluorobenzyl) derivative of deoxyribose by GC-MS. Fraction of new DNA was calculated based on the product/precursor relationship in samples from input and positive INTACT fractions. Bar graphs represent average DNA fraction new ± SEM, ** $p<0.01$ by two-tailed unpaired T test comparison (n=3/input and n=4/positive fraction).

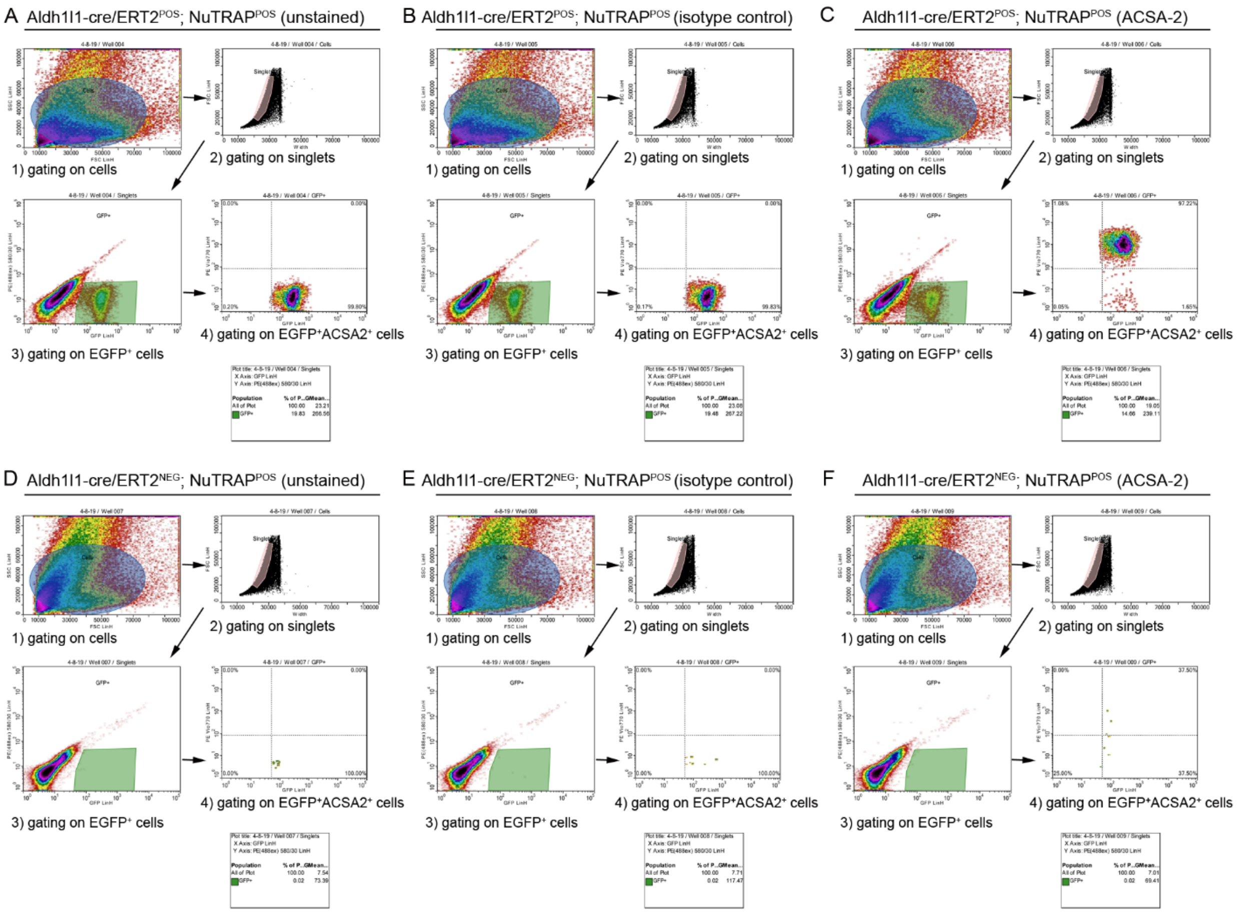


**Supplemental Figure 14. Gating strategy for flow cytometry analysis of tamoxifen induced cre recombination in the Aldh1l1-NuTRAP model.** Representative density plots from flow cytometry results illustrate steps followed for gating on EGFP$^+$ACSA2$^+$ cells (astrocytes) from single-cell suspension of Aldh1l1-cre/ERT2$^+$NuTRAP$^+$ brains analyzed. The gating process described under Methods section is applied to (A) unstained, (B) isotype control-PE-Vio770, and (C) ACSA-2-PE-Vio770 stained cells from the same brain cell suspension. 1) Cells are gated in the scatter range and for (2) subsequent selection of single cells (singlets). 3) Using a combination of filters, singlets are further gated for positive selection of EGFP$^+$ cells that are subsequently selected as ACSA2$^+$ (4: astrocytes). Same gating strategy applied to cre negative counterparts is represented in (D-E-F). Note: Samples in C and F are the same used to show representative data in Figure 1.

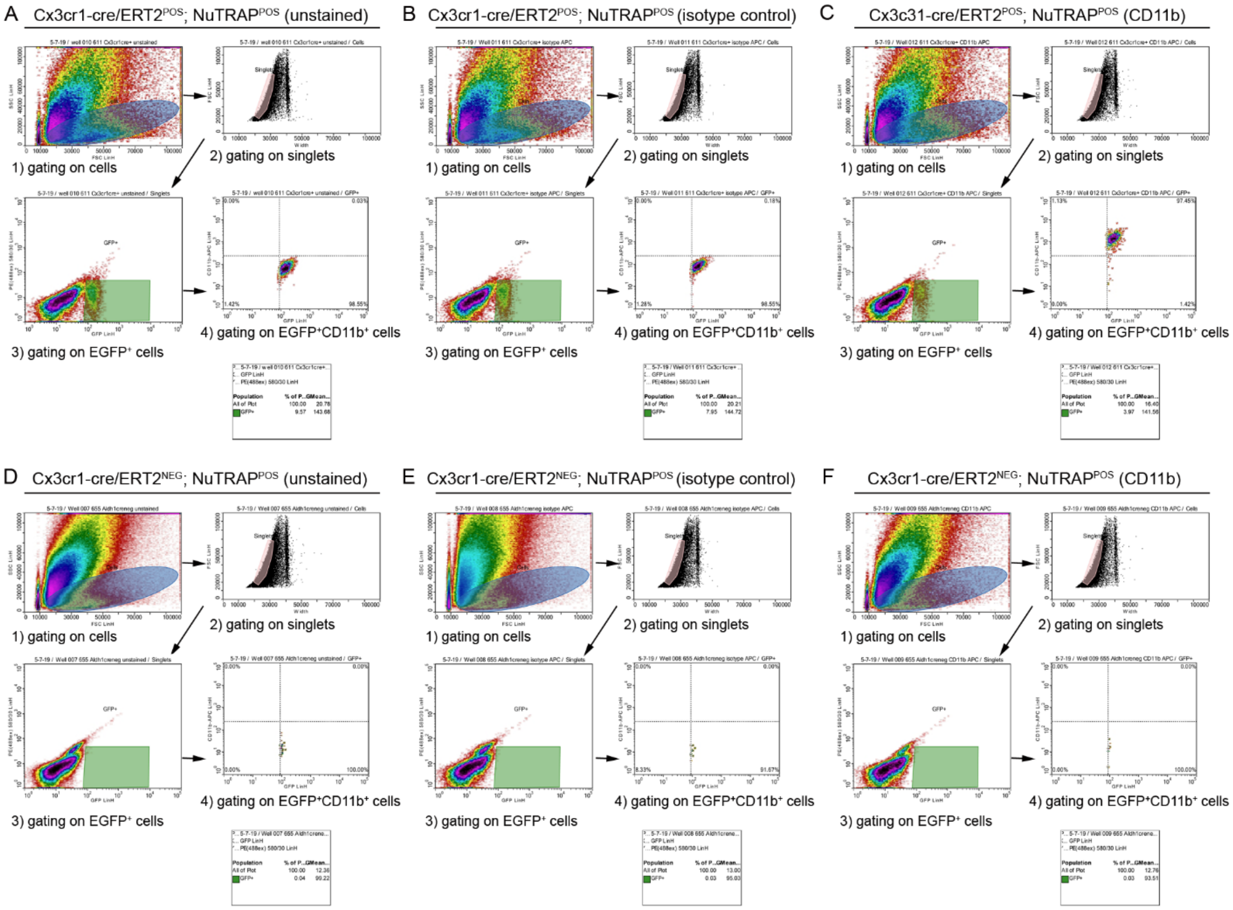


**Supplemental Figure 15. Gating strategy for flow cytometry analysis of microglia in the Cx3cr1-NuTRAP model.** Representative density plots from the flow cytometry results illustrate steps followed for gating on EGFP$^{+}$CD11b$^{+}$ cells (microglia) from single-cell suspension of Cx3cr1-NuTRAP brains analyzed. The gating process described under Methods section is applied to (A) unstained, (B) isotype control-APC, and (C) CD11b-APC stained cells from the same brain cell suspension. 1) Cells are gated in the scatter range for (2) subsequent selection of single cells (singlets). 3) Using a combination of filters, singlets are further gated for positive selection of EGFP$^{+}$ cells that are subsequently selected as CD11b$^{+}$ (4: microglia). Same gating strategy applied to cre negative sample (Aldh1l1cre/ERT2$^{-}$; NuTRAP$^{+}$) is represented in (D-E-F). Note: samples in C and F are the same used to show representative data in Figure 4.

| Type of product to amplify | primer ID (www.jax.org) | Description | expected product size |
|---|---|---|---|
| generic Cre (all Cre) | *OIMR 1084* | generic Cre Forward | |
| generic Cre (all Cre) | *OIMR 1085* | generic Cre Reverse | Transgene: ~100 bp |
| internal positive ctl | *OIMR 7338* | + ctl Forward | Internal positive CTL: 324 bp |
| internal positive ctl | *OIMR 7339* | +ctl Rev | |
| specific cre (microglia) | *20669* | Cx3cr1 Forward | Mutant: ~ 230 bp |
| specific cre (microglia) | *21058* | Cx3cr1 Common | Heterozigote: ~ 230 and 151 bp |
| specific cre (microglia) | *21059* | Cx3cr1 Mutant | Wild type: 151 bp |
| specific cre (astocytes) | *31091* | Aldh1l1 Tg Forward | |
| specific cre (astocytes) | *30308* | Aldh1l1 Tg Reverse | Transgene: 198 bp |
| internal positive ctl | *21238* | +ctl Forward | Internal positive ctl: 415 bp |
| internal positive ctl | *21239* | +ctl Reverse | |
| mCherry (floxed allele of NuTRAP) | *9703* | mCherry Forward | |
| mCherry (floxed allele of NuTRAP) | *9704* | mCherry Reverse | Transgene: 288 bp |
| internal positive ctl | *OIMR 8744* | internal positive ctl | Internal positive ctl: 200 bp |
| internal positive ctl | *OIMR 8745* | internal positive ctl | |
| NuTRAP(non floxed allele ) | *21306* | NuTRAP WT Forward | |
| NuTRAP(non floxed allele ) | *24493* | NuTRAP WT Reverse | Mutant: 194 bp |
| NuTRAP(floxed NuTRAP allele) | 32625 | NuTRAP Mutant Forward | Heterozygote: 109 bp and 194 bp |
| NuTRAP(floxed NuTRAP allele) | 32626 | NuTRAP Mutant Reverse | WT: 109 bp |

**Supplemental table 1. Primers used for genotyping of Aldh1l1- NuTRAP and Cx3cr1-NuTRAP mice (information available from Jax.org website).**

| Target gene for methyl primer | Target gene description | Methyl forward primer sequence 5 ► 3 | Methyl reverse primer sequence 5 ► 3 |
|---|---|---|---|
| Fabp7 | fatty acid binding protein 7, brain | TGATGTTTTGAGTAGGGATAGG | AAAAAAAAACAACTACAACCTCAA |
| Gpr84 | G protein-coupled receptor 84 | TTGTTTTAGAGAAAGGAAGGAA | TTCTAAAAACAACCAAACTTCC |
| Eno2 | enolase 2, gamma neuronal | GGTTAGAGTTAGGGGTTGGA | CTCCCCACCCTAATATCCTA |
| Aif1 | allograft inflammatory factor 1 | AGGTTTTTATTGTGTTGGTTTG | AATAACCATCCTACCTCAACCT |
| Ly96 | lymphocyte antigen 96 | TTTTAGTTGGGAATTGTTATTGA | CCAAATTAACCTTCTCCAAATA |
| Myd88 | myeloid differentiation primary response gene 88 | TGGGTTTTTTTTTTTAAAAAGT | AAACTACCCAACCTAAAAATCA |
| Tlr2 | Toll-like receptor 2 | GGGGTTTTTTTTATAATTGGAA | CCAACCTCCATACAAACCTAA |

**Supplemental table 2. Methyl primer sequences used for BSAS assays in the study.**

| gene/gene ID | Description | Taqman Gene Expression assay ID |
|---|---|---|
| *Aldh1l1* | Aldehyde Dehydrogenase 1 Family Member L1 | Mm03048957_m1 |
| *Gfap* | Glial Fibrillary Acidic Protein | Mm01253033_m1 |
| *Cx3cr1* | C-X3-C Motif Chemokine Receptor 1 | Mm00438354_m1 |
| *C1qa* | Complement C1q A Chain | Mm00432142_m1 |
| *Itgam* | Integrin Subunit Alpha M | Mm00434455_m1 |
| *Eno2* | Enolase 2 | Mm00469062_m1 |
| *Npas4* | Neuronal PAS Domain Protein 4 | Mm01227866_g1 |
| *Mog* | Myelin Oligodendrocyte Glycoprotein | Mm01279062_m1 |
| *Il1a* | Interleukin 1 Alpha | Mm00439620_m1 |
| *Il1b* | Interleukin 1 Beta | Mm00434228_m1 |
| *Tlr4* | Toll Like Receptor 4 | Mm00445274_m1 |
| *Myd88* | MYD88 Innate Immune Signal Transduction Adaptor | Mm01351743_g1 |
| *Tnf* | Tumor Necrosis Factor | Mm00443260_g1 |
| *Tlr2* | Toll Like Receptor 2 | Mm00442346_m1 |
| *Aif1* | Allograft Inflammatory Factor 1 | Mm00479862_g1 |
| *Gapdh* | Glyceraldehyde-3-Phosphate Dehydrogenase | 4352661 |

**Supplemental Table 3. Taqman gene expression assays used in qPCR analyses of the study.**

**Supplemental Table 4.** Gene lists from Figure 2.

**Supplemental Table 5.** Genes lists from figure 5.

**Supplemental Table 6.** Comprehensive information on the statistical analysis used in this study.

**Supplemental Table 7.** Instrument and settings (Immun
